# Glyceroglycolipids are essential for *Burkholderia cenocepacia* intracellular survival by preventing phagolysosome acidification

**DOI:** 10.1101/2023.05.30.542800

**Authors:** Holly Shropshire, Richard Guillonneau, Zengsheng Han, Rebekah A. Jones, Shadman Ahmed, Inmaculada García-Romero, Isabel Aberdeen, Ioannis Nezis, Miguel A. Valvano, David J. Scanlan, Yin Chen

## Abstract

*Burkholderia cenocepacia* is a problematic pathogen that infects people with cystic fibrosis and often causes fatal “cepacia syndrome”. *B. cenocepacia* infection is difficult to treat due to the high intrinsic resistance of the bacterium to antimicrobials and its ability to survive in macrophages. In this study, we uncover a hitherto unknown aspect of *B. cenocepacia*’s pathogenesis related to the formation of new glyceroglycolipids, which is required for intracellular survival. Using lipidomics, we observed that *B. cenocepacia* can produce three glyceroglycolipid species in phosphate deplete conditions using a PlcP-mediated lipid remodelling pathway originally discovered in soil and ocean-dwelling bacteria. While lipid remodelling as an adaptation strategy for environmental microbes to cope with the scarcity of phosphorus is known, its role in intracellular bacterial survival was not investigated. Using mammalian macrophages and *Galleria mellonella* larvae as infection models, we showed that the mutant unable to perform membrane lipid remodelling (Δ*plcP*) could not establish infection. Unlike the wild type bacterium, the Δ*plcP* mutant did not replicate within macrophages and failed to prevent phagosome acidification. Comparative genomics analyses showed that this PlcP pathway is conserved in all pathogenic *Burkholderia* that infect a variety of mammalian and plant hosts. Overall, our results indicate that membrane lipid remodelling plays an essential, yet previously overlooked, role in subverting host immunity.

## Introduction

*Burkholderia cenocepacia* is a member of the *Burkholderia cepacia* complex (Bcc), a group of at least 20 closely related *Burkholderia* species^1^. These are opportunistic human pathogens that typically infect people with cystic fibrosis and chronic granulomatous disease (CGD)^2, 3^. Many bacterial species can infect cystic fibrosis patients, the most prevalent one being *Pseudomonas aeruginosa* which has been extensively studied^4, 5^. Bcc infections are problematic due to the high intrinsic resistance of *Burkholderia* species to antimicrobials^6, 7^, the transmissibility of the infection across patients^8, 9^, and the poor prognosis after lung transplantation^10, 11^.

*B. cenocepacia* and other *Burkholderia* species can survive in immune cells such as macrophages^12, 13^, a common strategy for many intracellular bacterial pathogens^14^. Infected hosts have many countermeasures to restrict pathogens including withholding essential nutrients for microbial growth, a process known as nutritional immunity. This is best exemplified in the restriction of transition metals that are essential for microbial metabolism^15^. Recent evidence suggests that nutritional immunity can be extended to other essential microbial nutrients such as phosphorus^16, 17^. Microbial pathogens, on the other hand, have evolved sophisticated mechanisms to combat host restriction of essential nutrients. For example, *Salmonella* can adapt to phosphorus depletion during infection by upregulating the phosphorus starvation response, including upregulation of high affinity phosphate (Pi) uptake systems and utilisation of host-derived organic phosphorus to better cope with phosphorus limitation during host infection^14, 17^.

Membrane lipid remodelling is another bacterial strategy to cope with phosphorus limitation, widely studied in environmental microbes^18–21^, but not systematically investigated in pathogens. Bacterial membranes typically contain glycerophospholipids such as phosphatidylglycerol (PG) and phosphatidylethanolamine (PE). However, bacterial membrane lipids are much more complex and dynamic than previously thought^22^. We and others have shown that several microbes can substitute phosphorus-containing glycerophospholipids with surrogate non-phosphorus lipids to save the cellular phosphorus quota^18–25^. For example, this so-called PlcP-mediated lipid remodelling pathway allows cosmopolitan marine heterotrophic bacteria to be as competitive as their photosynthetic counterparts and to thrive in world’s most oligotrophic marine surface waters^19, 26^. In the legume-associated symbiotic bacterium, *Sinorhizobium meliloti*, a mutant unable to perform lipid remodelling grows poorly in phosphate limiting conditions, becoming less competitive in the natural environment^18^. Although the PlcP pathway is present in several major bacterial pathogens and our previous study on *Pseudomonas aeruginosa* revealed a role for lipid remodelling in antimicrobial resistance^21^, how lipid remodelling modulates infection is unknown. In this study, we investigate the link between lipid remodelling and virulence using the intracellular pathogen *B. cenocepacia* as the model system. We show lipid remodelling is required for *B. cenocepacia* virulence and plays a role in the intracellular survival of *B. cenocepacia* by preventing phagosome acidification.

## Results

### *B. cenocepacia* K56-2 produces glycolipids under phosphorus limitation

We have previously shown that *B. cenocepacia* K56-2 produces inorganic phosphate (Pi) scavenging proteins when grown under Pi-limited conditions^27^. Re-analysis of the previously published proteomics dataset identified three proteins encoded by a putative 3-gene operon, which were significantly upregulated under Pi-depletion^27^. This operon encoded a predicted PclP phospholipase (BCAL2364), a putative glycosyltransferase involved in glycoglycerolipid synthesis (BCAL2365), and a putative diacylglycerol kinase (BCAL2366, *dagK*) (**Figure 1A**). We hypothesised that this operon was involved in membrane lipid remodelling in response to Pi-deficiency in *Burkholderia cenocepacia* (**Figure 1A**).

**Figure 1.**
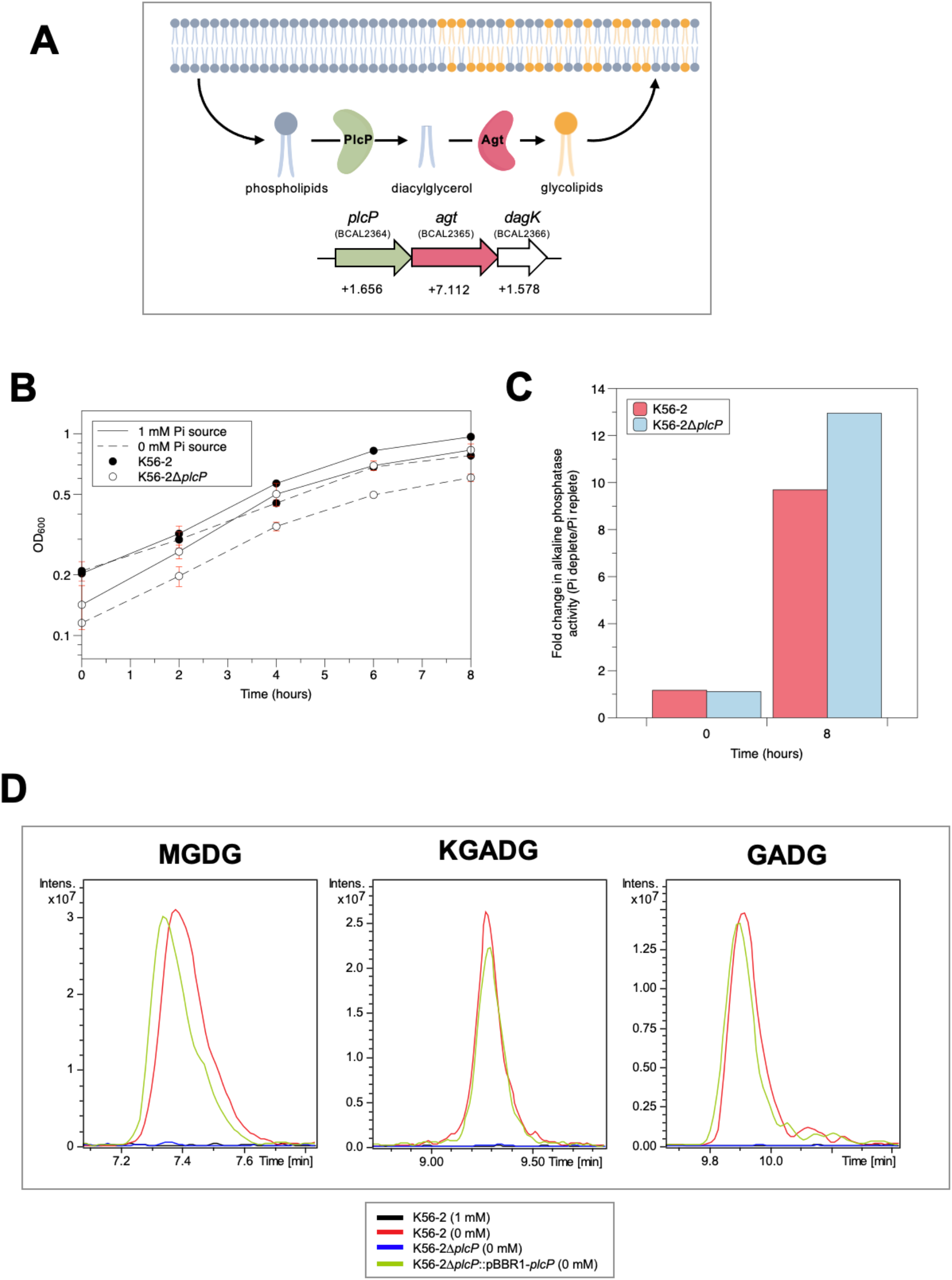
*B. cenocepacia* remodels its membrane lipids under phosphorus limitation. (A) The working model of key enzymes involved in lipid remodelling in *B. cenocepacia*. Lipid remodelling operon in *B. cenocepacia* consists of *plcP* (BCAL2364), *agt* (BPSL1190) and *dagK* (BCAL2366). Below the operon are the log2(fold change) values of these proteins in Pi-depletion compared to Pi-replete conditions unidentified in our previous comparative proteomics dataset^27^, showing that these proteins are significantly upregulated under Pi-limitation. (B) Growth curves of the wild type and the Δ*plcP* mutant grown under phosphate (Pi)-deplete (0 mM Pi source) and Pi-replete (1 mM Pi source) conditions in the defined MMA medium. Error bars represent ± standard deviation (n = 3). (C) Fold changes in alkaline phosphatase activity measured by quantifying the production of *p*-nitrophenol (*p*NP) from the hydrolysis of *p*-nitrophenol phosphate (*p*NPP). (D) Extracted ion chromatograms of the glycolipids MGDG, KGADG and GADG in positive ionisation mode (*m/z* 760, 772 and 814 respectively) which were extracted after 8 hours of growth in MMA. Black chromatogram is the wild type under Pi-repletion, red chromatogram is wild type under Pi-depletion, blue chromatogram is the Δ*plcP* mutant under Pi-depletion, green chromatogram is complementation mutant (K56-2Δ*plcP*::pBBR1-*plcP*) under Pi-depletion.

We investigated the membrane lipid profile of *B. cenocepacia* K56-2 when grown under Pi-limitation by performing comparative lipidomics via liquid chromatography-mass spectrometry (LC-MS) at the end of an 8-hour growth period in Pi-deplete (0 mM) and Pi-replete (1 mM) defined minimal medium (modified minimal A medium, MMA)^21^ (**Supplementary Fig. 1A**). LC-MS analysis identified three lipids that were present in the parental K56-2 under Pi-depletion but absent in the Pi-replete condition (**Figure 1D**). MS^n^ fragmentation (**Supplementary Figure 1B-D**) of the lipids eluting at ∼7.5 min, ∼9.5 min and ∼10 min denoted the presence of -glucosyl (neutral loss of 179), -ketoglucuronosyl (neutral loss of 191) and -glucuronosyl (neutral loss of 193) hydrophilic headgroups, respectively, suggesting that the three novel lipids are glycoglycerolipid monoglucosyl diacyglycerol (MGDG), ketoglucuronosyl diacylglycerol (KGADG) and glucuronosyl diacylglycerol (GADG)^19–21^ (**Figure 1D**). The production of these glycolipids required PlcP, since they were not found in the *plcP* deletion mutant, K56-2Δ*plcP*, but detected in the complemented mutant K56-2Δ*plcP* (pBBR1-*plcP*) (**Figure 1D**).

Moreover, K56-2Δ*plcP* and the wild type K56-2 strains grew at similar rates in the Pi-replete defined medium, but growth was delayed in K56-2Δ*plcP* under Pi depletion (**Figure 1B**). Under Pi limitation there was a 10- and 13-fold induction of alkaline phosphatase activity in K56-2 and K56-2Δ*plcP*, respectively (**Figure 1C**). These experiments support the notion that *B. cenocepacia* can remodel its membrane in response to Pi limitation by substituting membrane phospholipids with non-phosphorus containing glycoglycerolipids.

### PlcP is required for intracellular survival of *B. cenocepacia* in macrophages

*B. cenocepacia* is a facultative intracellular pathogen that survived in membrane vacuoles^13^. Since Pi can be a limiting nutrient for the survival and replication of bacterial pathogens within immune cells^17, 28^, we investigated whether Pi stress-induced lipid remodelling plays a role in the intracellular lifestyle of *B. cenocepacia*. For these experiments, a *plcP* deletion mutant was generated in strain MH1K, a gentamicin-sensitive derivative of *B. cenocepacia* K56-2, to enumerate the intracellular bacterial load using the gentamicin protection assay^29^. To facilitate the observation of *B. cenocepacia* intracellularly, mCherry was used to visualise *B. cenocepacia* strains. The murine macrophage cell line RAW264.7 and human macrophage cell line THP-1 were used, and these cells were cultured in a Pi-limited medium during the periods of interaction with the bacteria. Using confocal microscopy, we found that at 2 hours post infection (hpi) both WT and Δ*plcP* mutant were observed within macrophages, but at 24 hpi, Δ*plcP* appeared in much lower numbers (**Figure 2A** **and Supplementary Figure 2A**). Quantitative analysis by calculating the percentage of infected macrophages containing at least one bacterial cell shows the percentage of macrophages containing *B. cenocepacia* WT significantly increases over 24 hours of infection and nearly all macrophages were infected by the bacterium at 24 hpi. In contrast, the percentage of macrophages infected with the *plcP* mutant does not change significantly over time (**Figure 2B** **and Supplementary Figure 2B**). This suggests that the *plcP* mutant has a defect for intracellular survival, which was further supported by the analyses of intracellular bacterial load (colony forming unit, CFU) at each timepoint post-infection (**Figure 2C** **and Supplementary Figure 2C**). From 6 hpi to 24 hpi, there was a significant increase in bacterial cell numbers (∼ 2 log units) for the wild type, as previously described^29^, whereas the mutant cell numbers were significantly lower (*p* ≤ 0.05, *t*-test) (**Figure 2C** **and Supplementary Figure 2A**). Therefore, the lack of replication of the *plcP* mutant of *B. cenocepacia* in macrophages suggests that lipid remodelling is essential for invading macrophage immunity for successful replication and the long-term survival of *B. cenocepacia* intracellularly.

**Figure 2.**
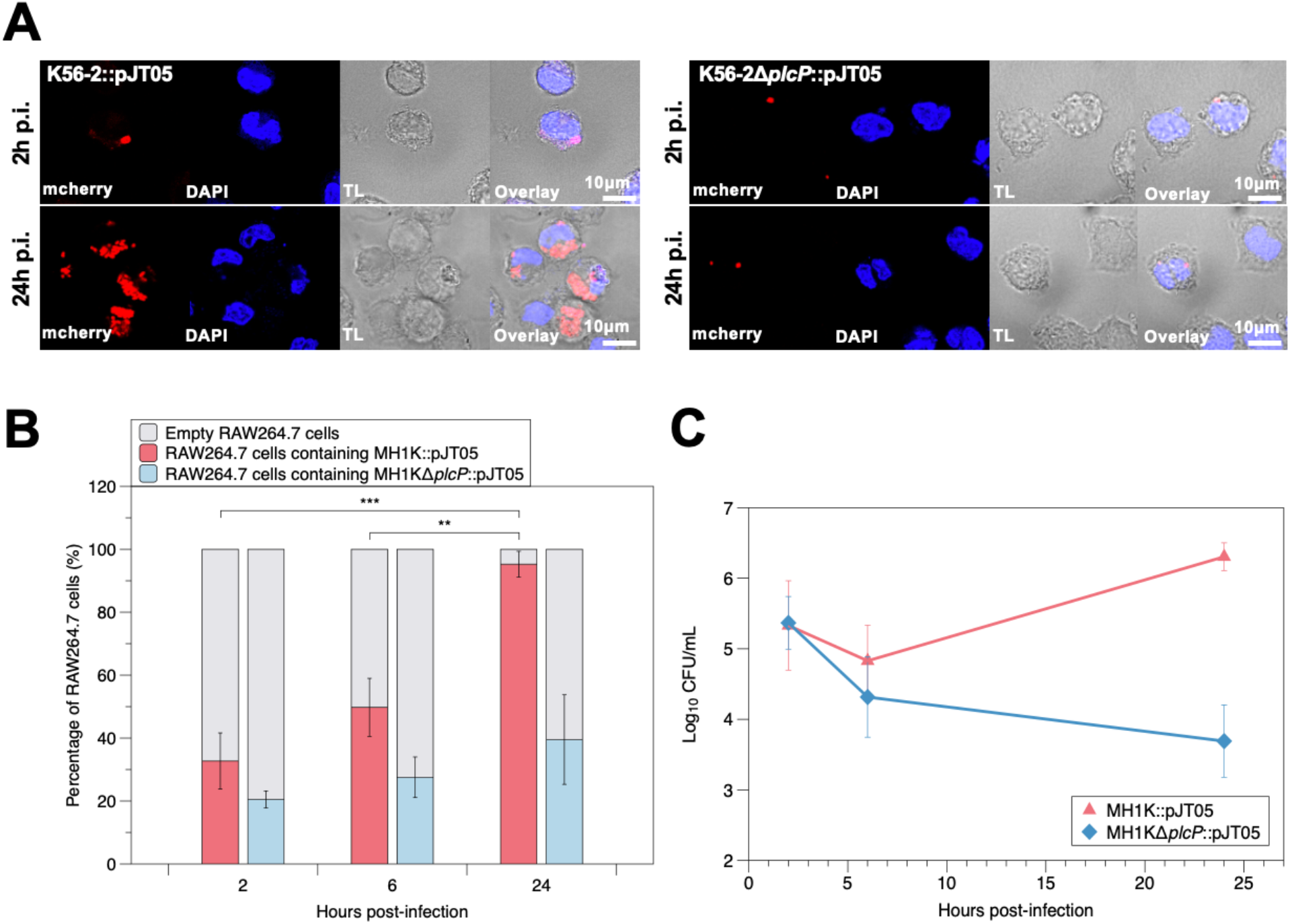
The *plcP* mutant of *B. cenocepacia* shows a defect for intracellular survival in RAW264.7 macrophages. (A) Confocal microscopy of the wild type (MH1K) and the *plcP* mutant (MH1KΔ*plcP*) containing mCherry expressing plasmid pJT05 in RAW264.7 macrophages at 2 and 24 hours post-infection (hpi). Red fluorescence indicates mCherry expressing bacterial cells, blue fluorescence indicates DAPI staining of the macrophage nucleus. TL: transmitted light microscopy image. Scale bars indicate 10 µm. Wild type bacteria replicate after 24 hours intracellularly contrary to the mutant which did not replicate intracellularly. (B) Percentage of RAW264.7 macrophages which contain at least 1 bacterial cell of the wild type (red) or the mutant (blue) expressing the mCherry plasmid, pJT05. Around 100 cells were counted in triplicate and error bars represent ± standard deviation. Significant increase in the wild type containing macrophages can be seen over time, but not for the mutant containing macrophages. ** *p* ≤ 0.01, *** *p* ≤ 0.001 (Independent t-test). (C) Bacterial survival measured as log_10_ CFU/mL recovered from RAW264.7 infected macrophages. Significant difference in CFU/mL between the wild type and the mutant at 24 hpi (*p* ≤ 0.01, independent *t*-test). Error bars represent ± standard deviation (n=3).

### PlcP is required for pathogenicity in the *Galleria mellonella* infection model

We investigated whether PlcP is required for *B. cenocepacia* virulence in an *in vivo* whole organism model using *Galleria mellonella*. Survival of infected larvae was measured over a 48-hour period post-infection after injection with either wild type K56-2 or K56-2Δ*plcP*. The percentage of wild type K56-2-infected surviving larvae decreased rapidly over 48 hours, and no viable larvae were found at 48 hours (**Figure 3A**). However, 78.3% of the Δ*plcP*-infected larvae remained alive at 48 hours (47 out of 60 larvae), confirming that the mutant has a significant defect to establish an infection in *Galleria mellonella* (logrank test *p*<0.001). A representative photograph of this experiment can be seen in **Figure 3B**, highlighting the clear difference in survival of infected larvae after 48 hours. Control experiments using larvae infected with heat-inactivated *B. cenocepacia* wild type or the mutant showed that all larvae survived across the 48-hour period (**Figure 3A**).

**Figure 3.**
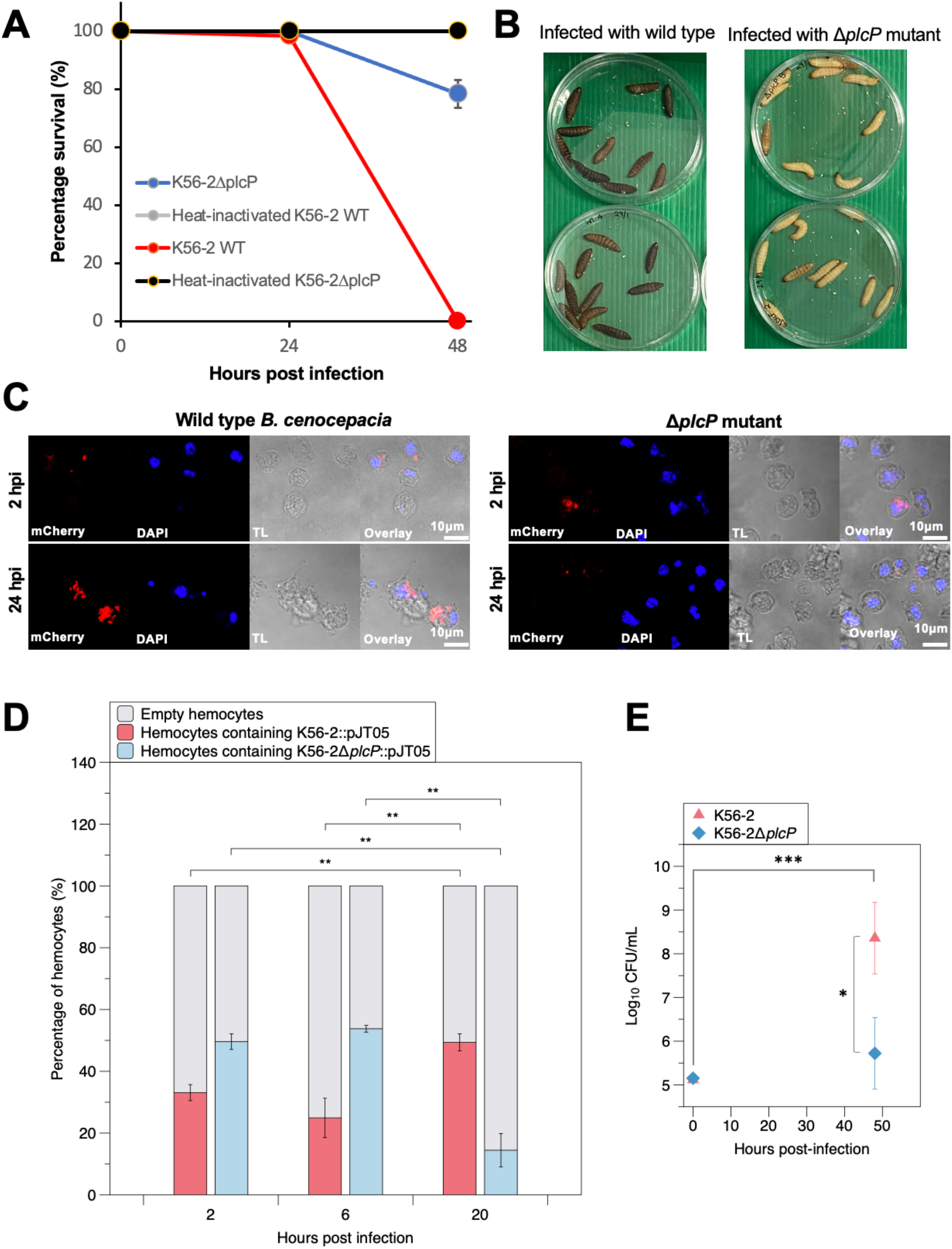
The *plcP* mutant of *B. cenocepacia* was unable to establish successful infection in the *Galleria mellonella* infection model. (A) *G. mellonella* larvae survival when infected with the wild type *B. cenocepacia* K56-2 (red) and the K56-2Δ*plcP* mutant (blue). Heat-inactivated K56-2 (grey) and K56-2Δ*plcP* (black) controls were also included and all larvae survived during the 48 hr experiment. Error bars represent ± standard deviation (6 biological replicates, 10 larvae per replicate). A logrank test showed significant difference of the survival rates of the insect larvae infected by the wild type K56-2 and the K56-2Δ*plcP* (*p* <0.001). (B) Photograph of experiment in Figure 3A showing two of the biological replicates at 48 hpi to show the stark contrast in melanization of *G. mellonella* larvae between wild type K56-2 and the K56-2Δ*plcP* mutant. (C) Confocal microscopy of the wild type and the *plcP* mutant containing mCherry expressing plasmid pJT05 in haemocytes at 2- and 24-hours post-infection (hpi). Red fluorescence indicates mCherry expressing bacterial cells, blue fluorescence indicates DAPI staining of the haemocytes nucleus. TL: transmitted light microscopy image. Scale bars indicate 10 µm. Wild type bacteria replicate after 24 hours intracellularly contrary to the mutant which did not replicate intracellularly. (D) Percentage of haemocytes which contain at least 1 bacterial cell of K56-2 (red) and K56-2Δ*plcP* (blue). ∼100 cells counted in triplicate. Error bars represent ± standard deviation. ** *p* ≤ 0.01, independent *t*-test. (E) Enumeration of wild type (red) and Δ*plcP* mutant (blue) from infected *G. mellonella* larvae at 0 and 48 hpi shown as log_10_ CFU/mL. Error bars represent ± standard deviation. * *p* ≤ 0.05, *** *p* ≤ 0.001, Kruskal-Wallis test.

To investigate whether bacterial survival was compromised during the infection inside of the insect, larvae were culled, and insect haemolymph was visualized under confocal microscopy. The results at 24 hpi show that wild type but not Δ*plcP* bacteria are present in haemocytes (**Figure 3C**), which was confirmed by counting the percentage of haemocytes-containing bacterial cells (**Figure 3D**). This is also supported by Transmission electron microscopy (TEM) imaging analysis which showed the presence of the wild type bacteria inside of *Burkholderia* containing vacuole in the insect haemocytes (**Supplementary figure 3**). Further, between 0 and 48 hpi, the numbers of wild type bacteria compared with Δ*plcP* increased significantly in the extracted haemolymph (**Figure 3E**), suggesting that the mutant is unable to survive or replicate in the larvae.

### PlcP is required to prevent phagosome acidification

Using a fluorescence-based reporter by fusing the *plcP* promoter region with a gene encoding mCherry, we followed *plcP* expression during *Galleria* haemocyte infection. These results showed that *plcP* transcription becomes readily detectable after 30 min post infection (**Figure 4A**), suggesting that the *plcP* operon is transcribed early after bacterial engulfment. One of the hallmarks of *B. cenocepacia* intramacrophage survival is the ability of the intracellular bacteria to survive in a late phagosomal vacuole that delays acidification^36^. Therefore, we investigated whether PlcP is required to prevent phagosome acidification using confocal microscopy of *G. mellonella* haemocytes infected with mCherry-labelled bacteria and stained with Lysotracker^TM^ Green. At 2 hpi, both the wild type and Δ*plcP* are associated with acidified vacuoles, but in contrast to the mutant, wild type bacteria inhibited phagolysosome acidification at 20 hpi (**Figure 4B**). Therefore, the Δ*plcP* mutant failed to both prevent phagolysosomal acidification and replicate intracellularly, becoming nearly cleared from the haemocytes (**Figure 4C**).

**Figure 4.**
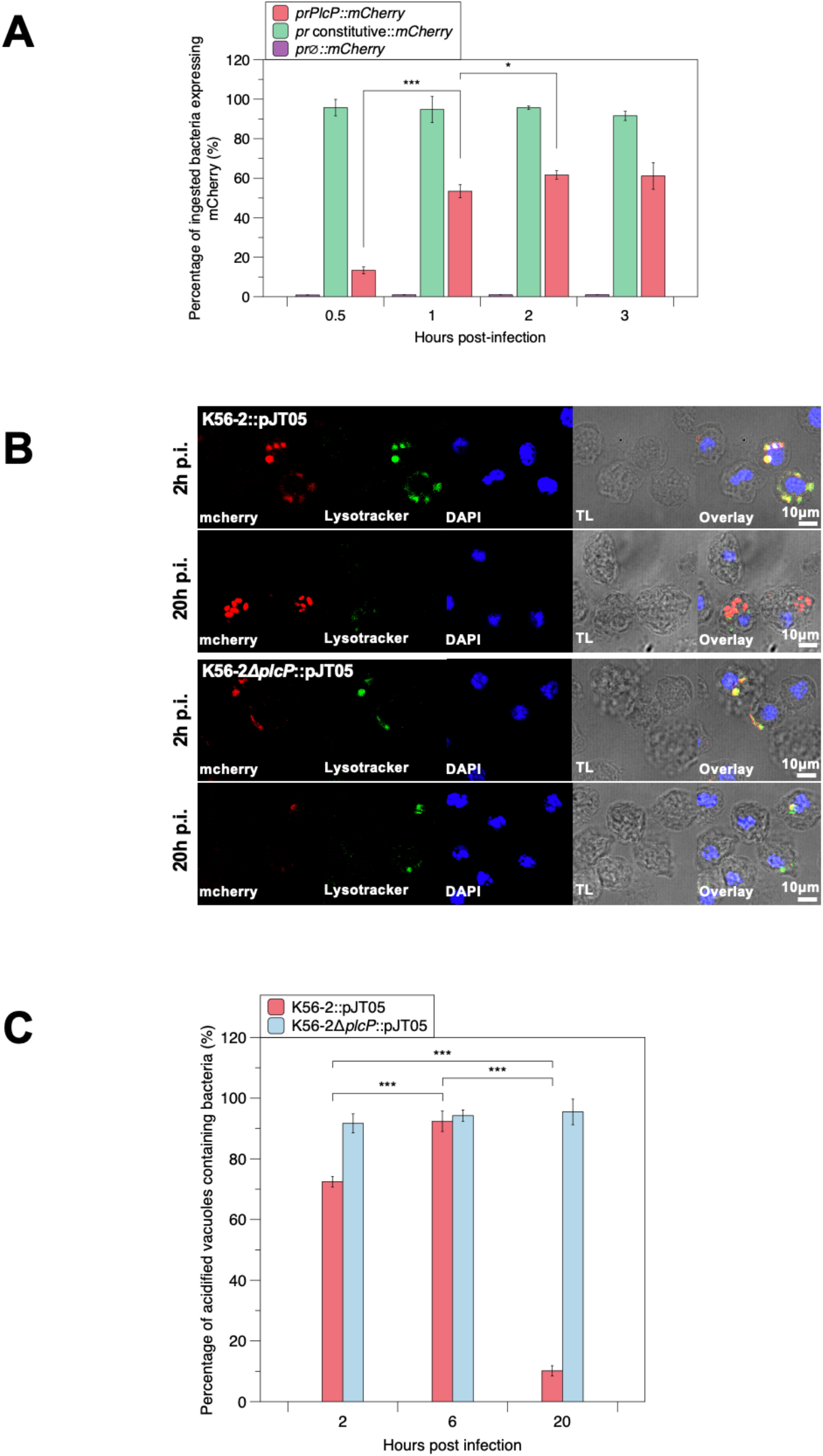
PlcP expression prevents phagolysosome acidification in *G. mellonella* haemocytes during infection. (A) Expression of PlcP in haemocytes using K56-2 with plasmids expressing mCherry from the putative *plcP* promoter (*prPlcP::mCherry)*, expressing mCherry constitutively (*pr* constitutive*::mCherry*) and mCherry with no promoter (*pr∅::mCherry*). ∼100 cells counted in triplicate. Error bars represent ± standard deviation. * *p* ≤ 0.05, *** *p* ≤ 0.001, independent *t*-test. (B) Confocal microscopy of K56-2 and K56-2Δ*plcP* containing mCherry expressing plasmid pJT05 in haemocytes at 2- and 20-hours post-infection (hpi). Red fluorescence indicates mCherry expressing bacterial cells, green fluorescence indicates acidification through Lysotracker green, blue fluorescence indicates DAPI staining of the macrophage nucleus. TL: transmitted light microscopy image. Scale bars indicate 10 µm. (C) Percentage of haemocytes containing K56-2 (red) and K56-2Δ*plcP* (blue) localized within an acidified compartment. ∼100 cells counted in triplicate. Error bars represent ± standard deviation. *** *p* ≤ 0.001, independent *t*-test.

### PlcP-mediated lipid remodelling pathway is conserved in pathogenic *Burkholderia* spp

To examine the conservation of the *plcP* pathway in *Burkholderia* genus, we searched the genome-sequenced *Burkholderia* strains (1302 genomes) deposited in the Integrated Microbial Genome and Microbiomes database at the Joint Genome Institute (JGI). The genus *Burkholderia* was first proposed in 1992, which was later split into *Burkholderia* spp. (pathogenic strains) and *Paraburkholderia* spp. (non-pathogenic environmental isolates) supported by comprehensive phylogenetics analyses^34, 35^. The phylogeny of *Burkholderia* spp. was reconstructed using the house keeping *rpoB* gene and the resulting phylogenetic tree classifies pathogenetic *Burkholderia* into three distinct clades (**Figure 5**). Clade Ia includes the Bcc strains, such as *B. cenocepacia*, *B. cepacia* and *B. multivorans.* Clade Ib includes *B. pseudomallei* and *B. mallei*, causative agents of human melioidosis and mammalian glanders respectively, whereas Clade Ic includes several phytopathogens (e.g., *B. glumae*, *B. gladioli* and *B. plantarii*) that infect various important crops^35^. All the genomes from species grouped into these three clades carried a predicted *plcP-agt-dagK* operon with a PlcP orthologue showing >84%, 91%, 72% amino acid sequence identity to the K56-2 PlcP, respectively (**Figure 5**). Thus, comparative genomics suggested that the ability to perform lipid remodelling through the PlcP pathway is conserved in pathogenic *Burkholderia* spp.

**Figure 5.**
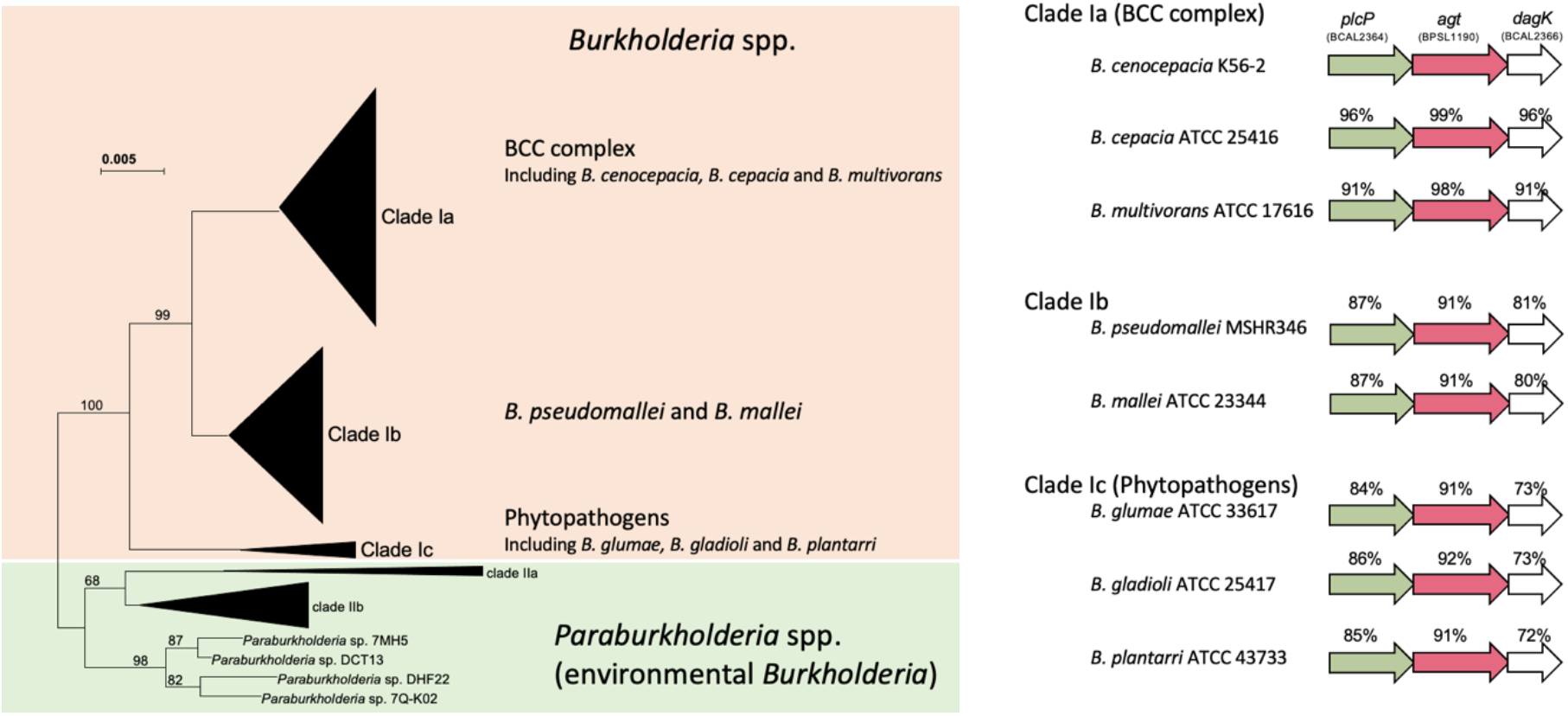
Comparative genomic analysis of the distribution of PlcP pathway in *Burkholderia* species, including human, animal and plant pathogens. Left, The evolutionary history of the housekeeping RpoB protein was inferred using the Neighbour-Joining method. RpoB from the newly established *Paraburkholderia* spp. (environmental *Burkholderia* isolates) were used as the outgroup. Clade Ia includes all Bcc complex bacteria; Clade Ib includes pathogenic *B. pseudomallei* and *B. mallei*; and Clade Ic includes phytopathogenic strains. The percentage of replicate trees in which the associated taxa clustered together in the bootstrap test (1000 replicates) are shown next to the branches. The evolutionary distances were computed using the Poisson correction method expressed as the number of amino acid substitutions per site. Evolutionary analyses were conducted in MEGA7. Right, Representative gene clusters of PlcP in human, animal and plant pathogenic *Burkholderia* spp. numbers above each gene show the sequence identity to corresponding gene in *B. cenocepacia* K56-2.

## Discussion

We report *B. cenocepacia* produces three types of glyceroglycolipids (MGDG, KDAGD, GADG) in response to phosphate stress in a *plcP-*dependent manner, suggesting that the PlcP enzyme plays a critical role to drive a membrane lipid remodelling in response to phosphate limitation. Our experiments also demonstrate that *plcP* is required for the intracellular survival of *B. cenocepacia* in mammalian macrophages and insect haemocytes. Using the *G. mellonella* infection model, we showed that during *B. cenocepacia* infection, PlcP transcriptional activity is expressed early after bacterial engulfment (**Figure 4A; Supplementary Figure 4**), suggesting that the *B. cenocepacia*-containing vacuole is a low Pi environment. This notion is consistent with the observation that Δ*plcP* cannot survive intracellularly (**Figure 2A**; **Figure 3C; Supplementary Figure 2A**). The role of PlcP in *B. cenocepacia* pathogenesis could be due to several possibilities. Bacterial stress in the *B. cenocepacia*-containing vacuole may not only be due to scarce concentrations of important ions (e.g. trace metals and phosphate) but also be attributed to increased production of oxidative reactive compounds. Intracellular *B. cenocepacia* delays phagosomal acidification and alters the assembly of the phagosomal NAPH oxidase in a Type VI secretion system (T6SS)-dependent manner by altering Rho family GTPases^36–37^. Through increasing the intracellular Pi pool by releasing phosphate from phospholipids, PlcP may also influence the regulation of the T6SS expression that is under the control of the global regulator AtsR^38^. However, it is likely that PlcP’s role in pathogenesis is multifaceted given that T6SS mutants are still able to survive intracellularly^39^.

The association of *plcP* with genes encoding putative lipid glycosyltransferase implicates that PlcP plays a key role in the formation of novel membrane lipids whereby phosphate head groups are removed by PlcP-type enzymes and replaced by other hydrophilic molecules (e.g., carbohydrates). This process of membrane lipid remodelling is widely known in environmental microbes as an adaptation strategy to cope with phosphorus limitation^18–20, 26^. However, a role for this process in bacterial infection had not been previously described. The lipidomics data from bacterial culture demonstrated the presence of novel glycolipids under Pi-deplete conditions which requires PlcP. Furthermore, promoter fusion assays clearly showed that the *plcP* is actively transcribed during infection, arguing for the role of lipid remodelling during infection although further experiment is required to unequivocally demonstrate the presence of such alternative surrogate lipids in the bacterial membrane during infection. However, our data support the notion that loss of membrane lipid remodelling in the absence of PlcP could explain the reduced survival of the mutant, hence their inability to manipulate the phagosome, in comparison with the wild type strain. Although the underlying basis for PlcP-mediated lipid remodelling in *B. cenocepacia*’s pathogenesis warrants further investigation, bacterial membrane lipids associated with virulence have been described before. Arguably the best studied example is sulfoglycolipids from *Mycobacterium tuberculosis*, the causative agent of tuberculosis. Biosynthesis of sulfoglycolipids is directly associated with intracellular survival of *M. tuberculosis* in human macrophages^30^ and the control of phagosomal acidification^32^. A recent study on the ornithine-containing aminolipid in *B. cenocepacia* J2315 showed that a mutant lacking expression of ornithine lipids was attenuated in the *G. mellonella* infection model^33^. Thus, manipulating membrane lipids may be an important yet understudied mechanism for pathogenicity.

Given that bacterial lipids are known to impact phagocytosis^30, 31^ as well as controlling phagosomal acidification^32^, we also sought to understand whether PlcP-mediated membrane remodelling of *B. cenocepacia* has an impact on the interaction between macrophages and the bacterium’s ability to survive stress. Phagolysosome represents a challenging environment for intracellular bacterial pathogens, typified with low pH, oxidative stress, and presence of an array of antimicrobials. However, there were no significant differences between the wild type and the *plcP* mutant when challenged by low pH, human serum nor by the addition of H_2_O_2_ or colistin (**Supplementary Figure 5**). Thus, it appears that the PlcP-mediated lipid remodelling is instrumental for the prevention of phagosome acidification (**Figure 4B**) as opposed to enhance the pathogen’s ability to resist environmental or antimicrobial insult associated with phagosomal maturation and acidification.

The results presented here uncover a new facet of understanding the success of this group of pathogens. Given that the PlcP pathway is conserved in all pathogenic *Burkholderia* species, membrane remodelling could be a widely used strategy by this group of bacteria to evade host immune cells. Our results support the notion that phosphorus is a key limiting nutrient for intracellular bacteria^14–17^ and its role in host-pathogen interactions should be further exploited. However, how does PlcP mediated lipid remodelling help prevent phagosomal acidification remains unclear. It is conceivable thought that PlcP-mediated membrane lipid remodelling is required for phagosomal acidification arrest^40–42^ or involved in the prevention of clearance through autophagy^43–44^ (**Figure 6**). Clearly the underlying molecular mechanisms underpinning the role of PlcP mediated lipid remodelling warrants further investigation. Nevertheless, this study reveals a new facet of phosphorus limitation induced membrane lipid renovation as a novel strategy for pathogenic bacteria to invade host immunity, thus arguing for the role of phosphorus limitation as an overlooked element of nutritional immunity.

**Figure 6.**
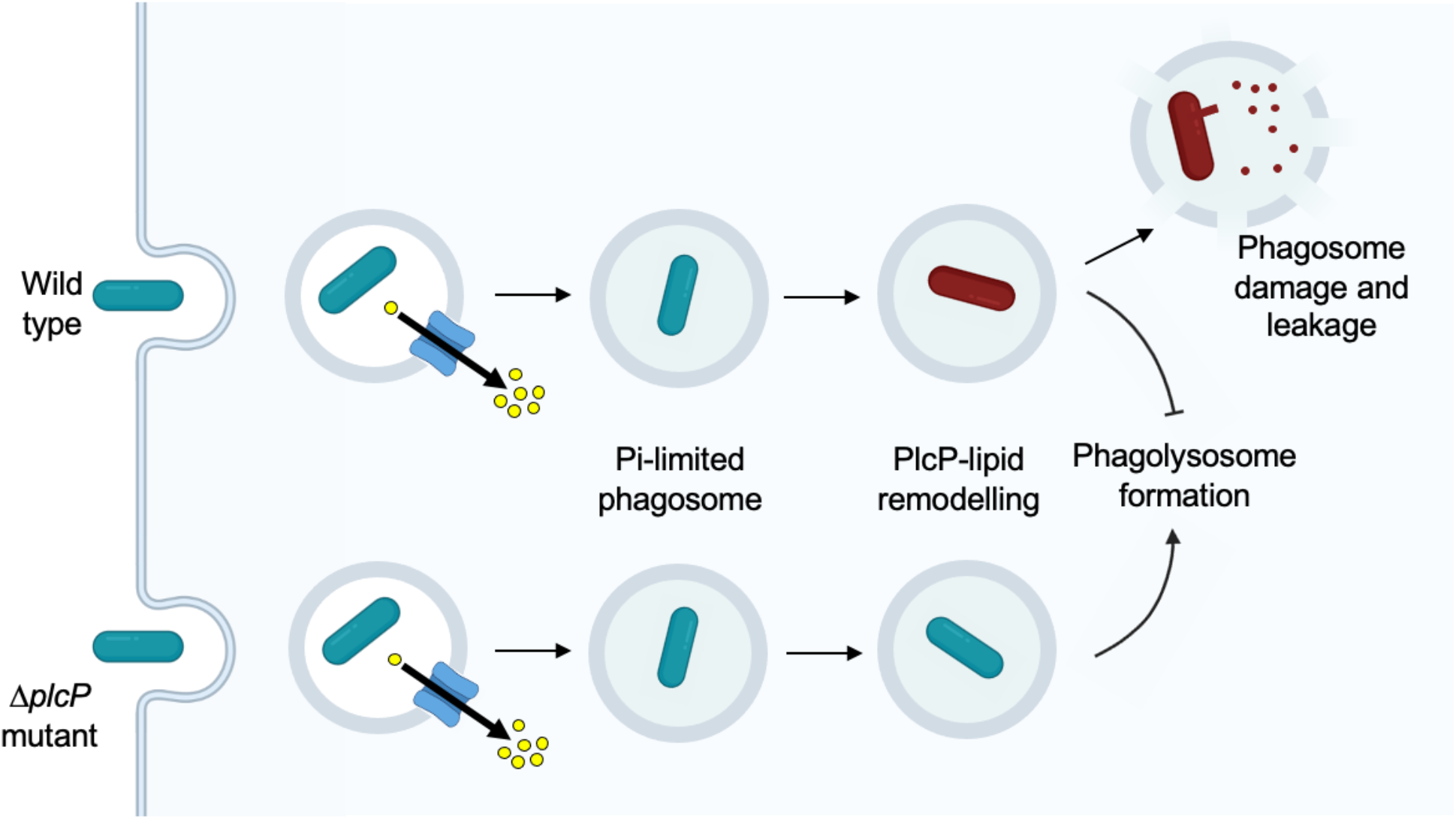
Hypothetical model of the role of PlcP-mediated lipid remodelling during *B. cenocepacia* infection. Firstly, *B. cenocepacia* is taken up by immune cells by phagocytosis. The host likely restricts the invasion of the bacterium by limiting essential nutrients within the phagosome, such as phosphate. In response to nutrient limitation, the bacterium swiftly remodels its membrane and induces the formation of several glycolipids and subsequent inhibits phagolysosome acidification. It is also likely that lipid remodeling allows the bacterium to subvert host autophagy pathways which also help to restrict bacterial infection.

## Methods

### Growth of *B. cenocepacia* and alkaline phosphatase activity assays

A single colony of the respective *B. cenocepacia* strain was inoculated into 50 mL LB media which was incubated overnight at 37°C at 180 rpm. The OD_600_ was measured, and culture was centrifuged for 10 mins at 4°C at 3100 x g (4000 rpm, Eppendorf 5810R benchtop centrifuge, rotor A-4-81). Pelleted cells were resuspended in the MMA medium^21, 27^ to 0.15 OD_600_. The high phosphate condition was also supplemented with 1 mM Na_2_HPO_4_. At each timepoint after inoculation, alkaline phosphatase activity was measured using *p*-nitrophenol phosphate (*p*NPP, final concentration 1 mM). Alkaline phosphatase activity was measured by quantifying *p*-nitrophenol (*p*NP) at 405 nm in a 96-well plate in triplicate using a FLUOstar Omega plate reader (BMG Labtech).

### Cultivation of macrophages

Mouse macrophage RAW264.7 and human macrophage THP-1 cell lines were obtained from ATCC and cultured with Dulbecco’s Modified Eagle Medium (DMEM) and Roswell Park Memorial Institute medium (RPMI 1640) respectively, both supplemented with 10% FBS and L-glutamine at 37 °C, 5% CO_2_. Low phosphate culture conditions were used during interaction with bacteria by using a modified Minimum Essential Medium (mMEM) with 10% FBS and L-glutamine containing 0.2 g/l CaCl_2_, 0.112 g/l MgSO_4_, 0.4 g/l KCl, 6.8 g/l NaCl, 1g/l Glucose, 5.958 g/l HEPES, 2.2 g/l NaHCO_3_, 20 ml/l of MEM AA solution 50X (Gibco^TM^, Waltham, MA, USA), 10 ml/l of MEM Non-Essential Amino Acids Solution 100X (Gibco^TM^, Waltham, MA, USA) and 10 ml/l MEM Vitamin Solution 100X (Gibco^TM^, Waltham, MA, USA).

### Construction and complementation of the *plcP* mutant in *Burkholderia cenocepacia* K56-2 and MH1K

The *plcP* gene (locus tag K56-2_17359, annotated as BCAL2364) was amplified by PCR and Gibson assembly was used to insert the *plcP* gene into the vector pGPI-SceI. Generation of *B. cenocepacia* deletion mutants was carried out as described previously^45^. Complementation of the *plcP* mutant was achieved by amplifying the *plcP* gene together with the upstream intergenic region using PCR and Gibson assembly to fuse this region into the broad-host-range cloning vector pBBR1MCS-3^46^. The plasmid was then introduced into the Δ*plcP* mutant through conjugation.

### Labelling *B. cenocepacia* using fluorescent proteins and construction of a fluorescent reporter

To visualize *B. cenocepacia* and its mutant within phagocytic cells, plasmid pJT05 containing the fluorescent protein gene *mCherry* was transformed into WT *B. cenocepacia* and *plcP* mutant by conjugation. To quantify the number of bacteria expressing *plcP* during *G. mellonella* and macrophage infections, we designed a reporter gene by fusing the native *plcP* promoter with *mCherry.* The fragments were firstly amplified by PCR and Gibson assembly was used to fuse *plcP* promoter with *mCherry* in the broad-host-range cloning vector pBBR1 MCS-3, hereinafter named *prPlcP::mCherry*. Controls containing only *mCherry* without a promoter (mCherry not expressed) and *mCherry* with the 200 bp upstream of *mCherry* in pJT05 (constitutively expressed) were designed by the same method, hereafter named *pr ∅::mCherry* and *pr* constitutive*::mCherry*, respectively. These plasmids were transformed into *Escherichia coli* WM3064 via electroporation and mobilized into WT *B. cenocepacia* K56-2 via conjugation. Briefly, overnight culture of WM3064 transformed with the desired plasmids was mated with overnight culture of *B. cenocepacia* in a 3:1 ratio on SOB plates containing 100 μg/mL of 2,6-diaminopimelic acid (DAP, Sigma-Aldrich, St. Louis, MO, United States) for 2 days at 37°C. *B. cenocepacia* transconjugants were selected on DAP-free LB agar plates containing 150 μg/mL of tetracycline. In order to visualize bacteria not expressing mCherry, prior to infection experiment *B. cenocepacia* cells was stained with SybrGreen at 1X during 20 min.

### *B. cenocepacia* infection of macrophage cell lines

Prior to infection, WT *B. cenocepacia* and the *plcP* mutant strains were cultured in MMA broth until an OD_600nm_ of 0.6. To initiate infection of macrophages (RAW264.7 and THP-1), bacteria and macrophages were transferred to mMEM medium. Macrophage suspensions were inoculated into 24-well microplates at 5 x 10^5^ cells/ml containing a coverslip previously UV-sterilized and incubated overnight at 37 °C, 5% CO_2_ to initiate macrophages attachment. The bacteria (WT *B.* cenocepacia MH1K or the MH1K *plcP* mutant, both containing pJT05 constitutively expressing mCherry) were inoculated at a multiplicity of infection (MOI) of 2 (equivalent to 2 bacteria per macrophage cells). Low speed centrifugation (5 min at 900 × g) was used to slowly move the bacteria down toward the adhered macrophages, and to initiate and synchronize physical interaction between bacteria and macrophages. The microplate was then incubated at 37 °C, 5% CO_2_. After 2h post infection (hpi), the cells were washed three times with mMEM to remove extracellular bacteria and a gentamicin protection assay (200 μg/mL) was performed for 2h to kill any residual extracellular bacteria for the time points 6h and 24h. After 2, 6, 24 hpi, bacteria-macrophage interaction subsamples were stained with DAPI (5 μg/mL). Afterward, samples were directly fixed by adding formaldehyde (4% [vol/vol]) for 30 min. Finally, cells were mounted with a drop of Mowiol antifade before observation using confocal laser scanning microscopy (CLSM, Zeiss LSM 880, Göttingen, Germany).

To quantify bacterial load inside macrophages, macrophages suspended in mMEM were inoculated into 96-well microplates at 5 x 10^5^ cells/ml and incubated overnight at 37 °C, 5% CO_2_ to initiate macrophages attachment. The bacteria (WT and Δ*plcP*) were inoculated at a MOI of 2 into 96-well microplates containing macrophages and low speed centrifugation was used to initiate and synchronize physical interaction between bacteria and macrophages. The microplates were then incubated at 37 °C, 5% CO_2_. After 2 hpi, the cells were washed three times with mMEM to remove extracellular bacteria and gentamicin protection assay (200 μg/mL) was performed for 2h at 6 and 24 hpi. At each times point, macrophages were treated with Triton X-100 (0.05%, w/v) for 20 min at room temperature and mechanically lysed by mixing using a syringe and 25-g needle (BD, Franklin Lakes, NJ, USA), as previously described^47^. Bacteria were then enumerated by serial dilution on LB agar plates.

### Infection of *Galleria* haemocytes by *B. cenocepacia*

Galleria mellonella larvae (LiveFood, Rooks Bridge, UK) weighing 0.25-0.3 g and with no pigmentation were selected for experimentation within a 10-day period from their arrival (stored at 4°C). Prior to infection, WT K56-2 *B. cenocepacia* and the K56-2 *plcP* mutant strains were cultured in LB broth until an OD_600nm_ of 0.6. Bacterial cultures were resuspended in PBS 1X and a serial dilution was performed to reach a concentration of 20,000 bacterial cell/mL. A 500-μl Hamilton syringe equipped with an automatic dispenser was used to inject 10 μl of bacterial suspension into *G. mellonella* via the hindmost left proleg as previously described^48^.

In order to assess the survival of the worms, 10 larvae per biological replicate were incubated in sterile petri dishes after infection for 48 hours at 37°C and survival was monitored every 24 hours by response to touch. One set of larvae was injected with sterile PBS 1X and one set of larvae was not injected, as controls to ensure the worms’ health.

To estimate the survival of bacteria inside the larvae, haemolymph from all 10 larvae in each biological replicate were culled after 48 hpi. Immediately after collection, haemolymph was serially diluted and 10 μL was plated onto LB containing 100 μg/mL gentamycin (Sigma-Aldrich, Saint Louis, MO, USA) and 200 μg/mL ampicillin (Sigma-Aldrich, Saint Louis, MO, USA) to kill any natural gut microbiota using the Miles and Misra method^49^. Initial dose of bacteria was calculated by serially diluting the culture used for injection and plating 100 μL onto LB agar plates.

In order to observe the fate of the bacteria in the haemocytes of the inset, 5 larvae per sample were injected, alongside uninfected and PBS-injected controls. After 2, 6, 20 hpi, injected larvae were culled and the haemolymph was extracted and resuspended in 1 ml of insect physiological saline (IPS) which consisted of; 150 mM NaCl, 5 mM KCl, 30 mM trisodium citrate, 10 mM EDTA, 100 mM Tris/HCl (pH 6.9) in sterile water; in a 24-well microplate containing a UV-sterilised glass coverslip. The haemocyte samples were then incubated for 1h at 37°C to initiate haemocytes attachment to coverslips. After incubation, the cells were washed three times with IPS to remove unattached haemocytes before staining with 70 nM Lysotracker Green (Thermofisher, Waltham, MA, USA) and 5 μg/mL DAPI (Sigma-Aldrich, Saint Louis, MO, USA) for 30 min at 37°C, 60 rpm. Afterward, samples were directly fixed by adding formaldehyde (4% [vol/vol]) for 30 min. Finally, cells were mounted with a drop of Mowiol antifade before observation using CLSM.

To visualise bacterial inside of the insect haemocytes by transmission electron microscopy (TEM), insect haemolymph was extracted after 20 hpi of the wild type K56-2 infected larvae (n=8), washed with IPS three times before fixing with 1 mL PBS containing 2.5% (w/v) glutaraldehyde. After 3 washes with PBS, the plates were then stored at 4°C before sample preparation which was completed by The Advanced Bioimaging RTP (The University of Warwick). Briefly, TEM samples were postfixed by immersion in 1% osmium tetroxide in water for 1 hour, followed by staining with 2% uranyl acetate for 1 hour. Samples were dehydrated using increasing acetone concentrations and directly embedded from the surface into Low Viscosity resin (Agar Scientific, UK). Ultrathin (70 nm) sections were stained with uranyl acetate followed by lead citrate. Samples were imaged using a Jeol 2100Plus transmission electron microscope (Jeol, Tokyo, Japan) operating at 200 kV, equipped with a Gatan OneView IS detector (Gatan Ametek, USA).

### Lipidomics analysis using LC-MS

A single colony of the respective *B. cenocepacia* strain was inoculated into the MMA medium to OD_600_ of 0.15. Cells were grown for 8 hours in the respective Pi-condition. Lipids were extracted using a modified Folch extraction method ^50^. A volume of cells equals to 1 mL of 0.5 OD_600_ was centrifuged for 15 mins at 4°C at 3100 x g. The supernatant was discarded and 50 μL lipid standard (10 μM), sphingosyl PE (SPE), was added giving a final concentration of 500 nM (Avanti Lipids). The pellet was then resuspended in 500 μL ice-cold methanol by vortexing. 300 μL ice-cold sterile water was added to the cells, followed by 1 mL ice-cold chloroform. The sample was vortexed for 30 seconds before being centrifuged for 15 mins at 4°C at 1739 x g (3000 rpm, Eppendorf 5810R benchtop centrifuge, rotor A-4-81). The lower phase containing the lipids was transferred to a fresh glass vial and the solvent was removed using a stream of nitrogen gas. Dried lipids were resuspended in 1 mL of acetonitrile:ammonium acetate (95:5, vol/vol). Samples were then analysed by LC-MS using a Dionex Ultimate 3000-LC/MS (amaZon SL, Bruker). Lipids were separated based on the polarity of their head group on a BEH Amide XP column (Waters). The column was maintained at 30°C with a flow rate of 150 μL/min. Each run consisted of a 10-minute period of equilibration with 95% acetonitrile and 5% 10 mM ammonium acetate (pH 9.2). After this, 5 μL sample was injected into the system and was run on a 15 min gradient of 5 to 30% 10 mM ammonium acetate (pH 9.2). Each sample was analysed in positive and negative ionization modes with auto fragmentation.

### Sensitivity assays

To test whether WT *B. cenocepacia* and the *plcP* mutant exhibit the same resistance to various stresses, sensitivity assays against human serum (Merck Millipore, Burlington, MA, USA), colistin (Sigma-Aldrich, Saint Louis, MO, USA), pH and H2O2 (Fisher, Waltham, MA, USA) were performed. Overnight cultures of *B. cenocepacia* in MMA were diluted in fresh medium to an OD_600nm_ of 0.0055 containing either: 0%, 20% [vol/vol] or 30% [vol/vol] of serum or 0 μg/mL, 256 μg/mL or 512 μg/mL of colistin sulfate salt, or 0 mM or 1.5 mM of H_2_O_2_ for serum, colistin and H_2_O_2_ assay respectively. To test the pH sensitivity, overnight cultures of bacteria in MMA were diluted to an OD_600nm_ of 0.0055 in fresh medium adjusted to pH 7 or 5.5 with 10 mM MES buffer (Sigma-Aldrich, Saint Louis, MO, USA). The different sensitivity assays were then incubated at 37°C, 180 rpm, for 0h and 2h for the serum, H_2_O_2_ and pH assay and for 0h and 4h for the colistin assay. At each time point, bacterial concentrations were quantified by serial dilution on LB agar plates.

### Statistics

For each experiment, at least three biological replicates were performed and ≥ 100 macrophages or haemocytes per replicates were counted. To test for statistically significant differences (*p* < 0.05) between two or more conditions, a *t* test or a Kruskal-Wallis test was performed. To compare the survival rate of *G. mellonella* infection by wild type *B. cenocepacia* K56-2 and its *plcP* mutant, a logrank test was performed. These tests were performed by using SPSS 26.0 (IBM, Armonk, NY, USA).

## Acknowledgement

We acknowledge funding from the BBSRC and MIBTP (BB/M01116X/1) doctoral training centre to H.S and I.A, the MRC doctoral training centre to R.A.J, the China Scholarship Council to Z. H. (202007815003) and European Research Council (grant agreement no. 726116) to Y.C. We thank Saskia Bakker at the University of Warwick Advanced Bioimaging Research Technology Platform for help with transmission electron microscopy.

## Notes

### Competing Interest Statement

The authors have declared no competing interest.

