## Supplementary files for "Glyceroglycolipids are essential for *Burkholderia cenocepacia* intracellular survival by preventing phagolysosome acidification"

**Supplementary Information**

**Supplementary Figure 1.** LC-MS analysis of *B. cenocepacia* membrane lipids.

**Supplementary Figure 2.** The *B. cenocepacia plcP* mutant shows a defect in intracellular survival in human THP-1 macrophages.

**Supplementary Figure 3.** Transmission electron microscopy (TEM) image of wild type K56-2 infected insect haemocyte at 20 hour post infection (hpi).

**Supplementary Figure 4.** Active transcription of the *B. cenocepacia plcP* gene in wild type cells during infection of *Galleria mellonella* using a promoter fusion assay.

**Supplementary Figure 5.** Sensitivity of wild type *B. cenocepacia* and the *plcP* mutant to various stresses.

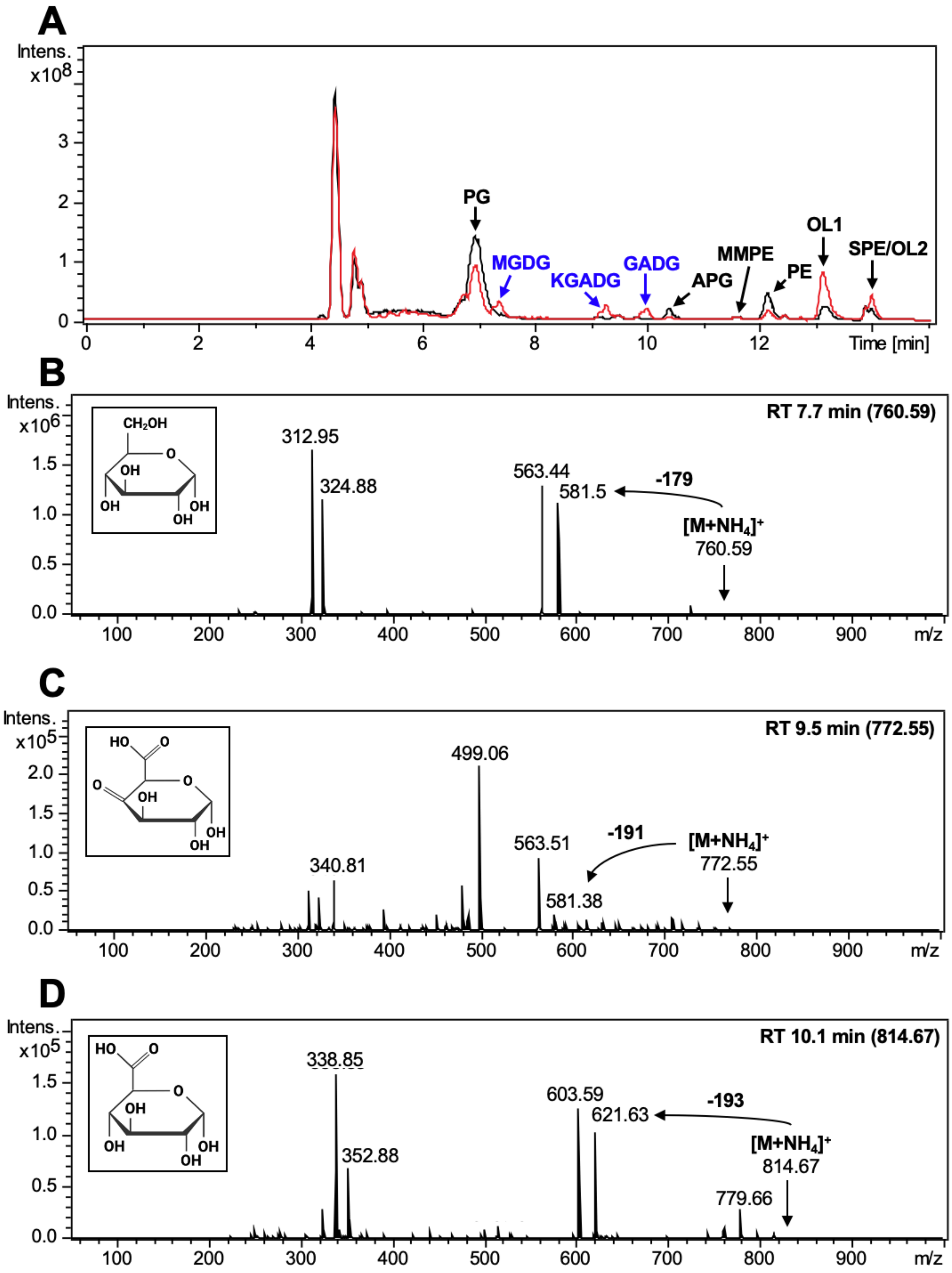
A
Intens. x10^8
PG
MGDG
KGADG
GADG
APG
MMPE
PE
OL1
SPE/OL2
Time [min]
B
Intens. x10^6
RT 7.7 min (760.59)
CH2OH
312.95
324.88
563.44
581.5
-179
[M+NH4]+
760.59
m/z
C
Intens. x10^5
RT 9.5 min (772.55)
499.06
563.51
-191
[M+NH4]+
772.55
340.81
581.38
m/z
D
Intens. x10^5
RT 10.1 min (814.67)
338.85
603.59
621.63
-193
[M+NH4]+
814.67
352.88
779.66
m/z

**Supplementary Figure 1. LC-MS analysis of *Burkholderia cenocepacia* membrane lipids.** (A) Full chromatogram, in positive ionisation mode, of the *B. cenocepacia* lipidome under Pi-deplete (red) and Pi-replete (black) growth conditions. Lipids are indicated as follows: PG, phosphatidylglycerol; MGDG, monoglucosyl diacylglycerol; KGADG, keto-glucuronic acid diacylglycerol; GADG, glucuronic acid diacylglycerol; APG, alanyl-phosphatidylglycerol; MMPE, monomethylated phosphatidylethanolamine; PE, phosphatidylethanolamine; OL1, ornithine lipid type 1; SPE, sphingosyl-phosphatidylethanolamine (internal standard); OL2, ornithine lipid type 2.

(B) $MS^2$ fragmentation of the MGDG peak at 7.7 min with parent ion 760.59. A neutral loss of 179 *m/z* corresponds to the loss of the glucosyl head group of MGDG.

(C) $MS^2$ fragmentation of the KGADG peak at 9.5 min with parent ion 772.55. A neutral loss of *191* m/z corresponds to the loss of the keto-glucuronosyl head group of KGADG.

(D) $MS^2$ fragmentation of the GADG peak at 10.1 min with parent ion 814.67. A neutral loss of 193 *m/z* corresponds to the loss of the glucuronosyl head group of GADG. All fragmentations are shown in positive ionisation mode.

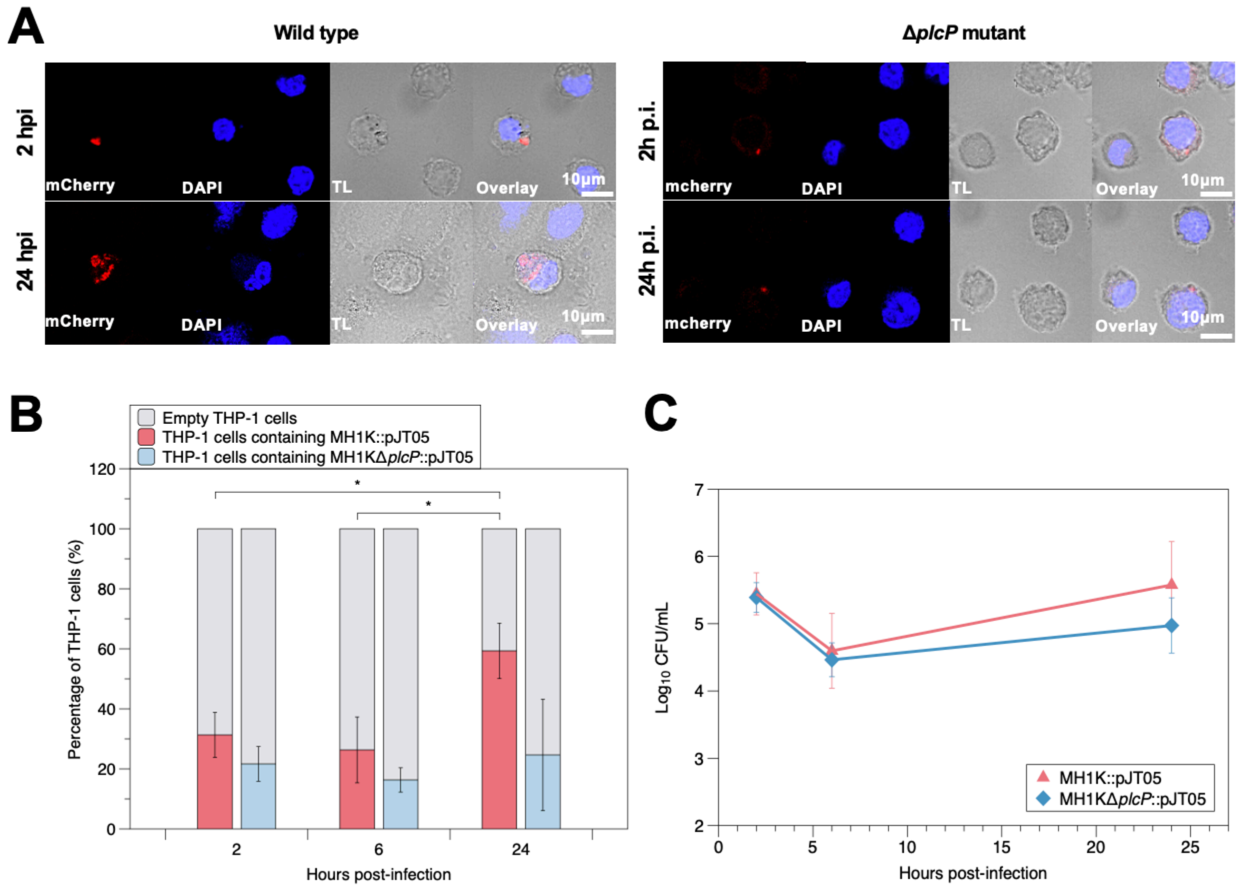


**Supplementary Figure 2. The *B. cenocepacia plcP* mutant shows a defect in intracellular survival in human THP-1 macrophages.** (A) Confocal microscopy of the wild type and the *plcP* mutant containing the mCherry expressing plasmid pJT05 in THP-1 macrophages at 2 and 24 hours post-infection. Red fluorescence indicates mCherry-expressing bacterial cells, blue fluorescence indicates DAPI staining of the macrophage nucleus. TL: transmitted light microscopy image. The scale bar is 10 µm. (B) The percentage of THP-1 macrophages which contain at least one bacterial cell in the wild type (red) or the *plcP* mutant (blue). ~100 cells were counted in triplicate and error bars represent the mean ± standard deviation. A significant increase in *B. cenocepacia* cells in wild type-infected macrophages can be seen over time, but not for the mutant-infected macrophages. *= $p \leq 0.05$ (Independent t-test, SPSS statistics). (C) Bacterial survival measured as log10 CFU/mL recovered from infected THP-1 macrophages. Error bars represent mean ± standard deviation (n=3).

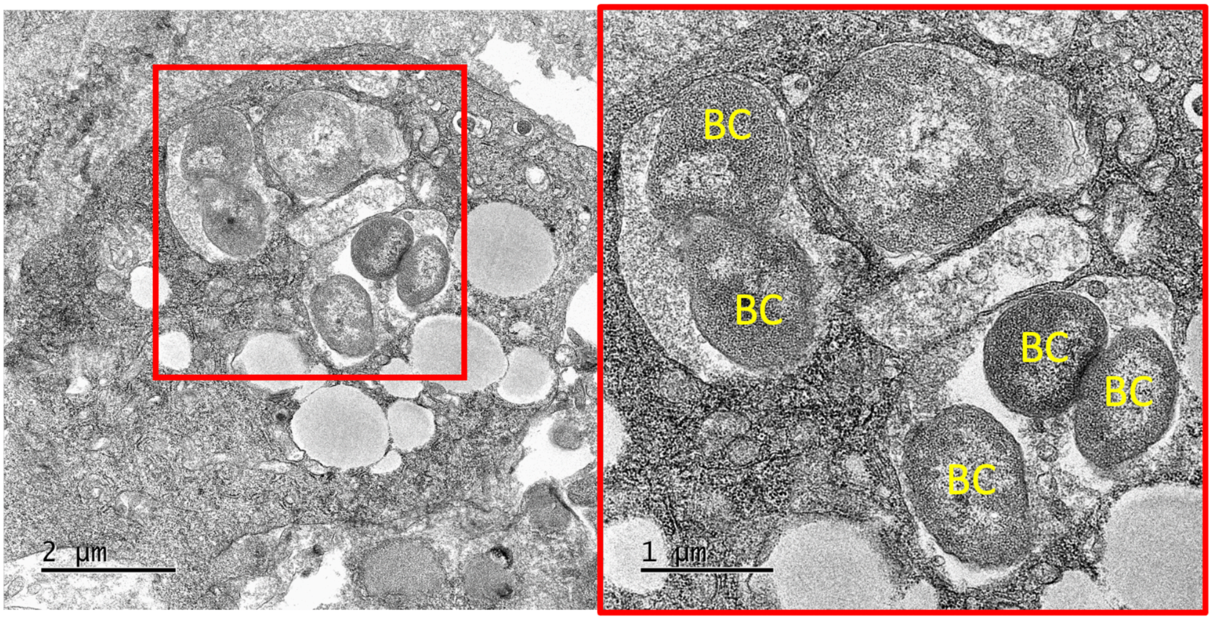


**Supplementary Figure 3. Transmission electron microscopy (TEM) image of wild type K56-2 infected insect haemocyte at 20 hour post infection (hpi).** Right-hand side panel shows zoomed in image of the haemocyte (red box of left-hand side panel). Bacterial cells are labelled BC.

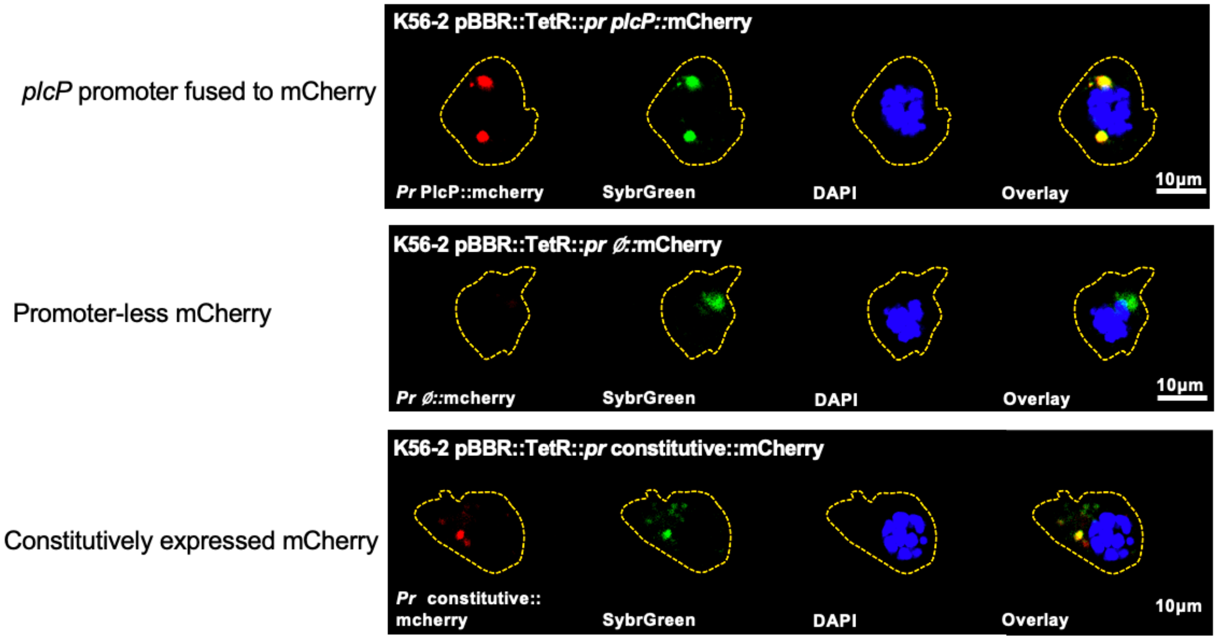


**Supplementary Figure 4. Active transcription of the *B. cenocepacia plcP* gene in wild type cells during infection of *Galleria mellonella* using a promoter fusion assay.** Top panel, mCherry fused with the *plcP* promoter sequence. Middle panel, control plasmid of mCherry without promoter. Bottom panel, control plasmid of constitutively expressed mCherry. Bacterial cells are stained with SybrGreen and the nucleus of haemocytes is stained with DAPI.

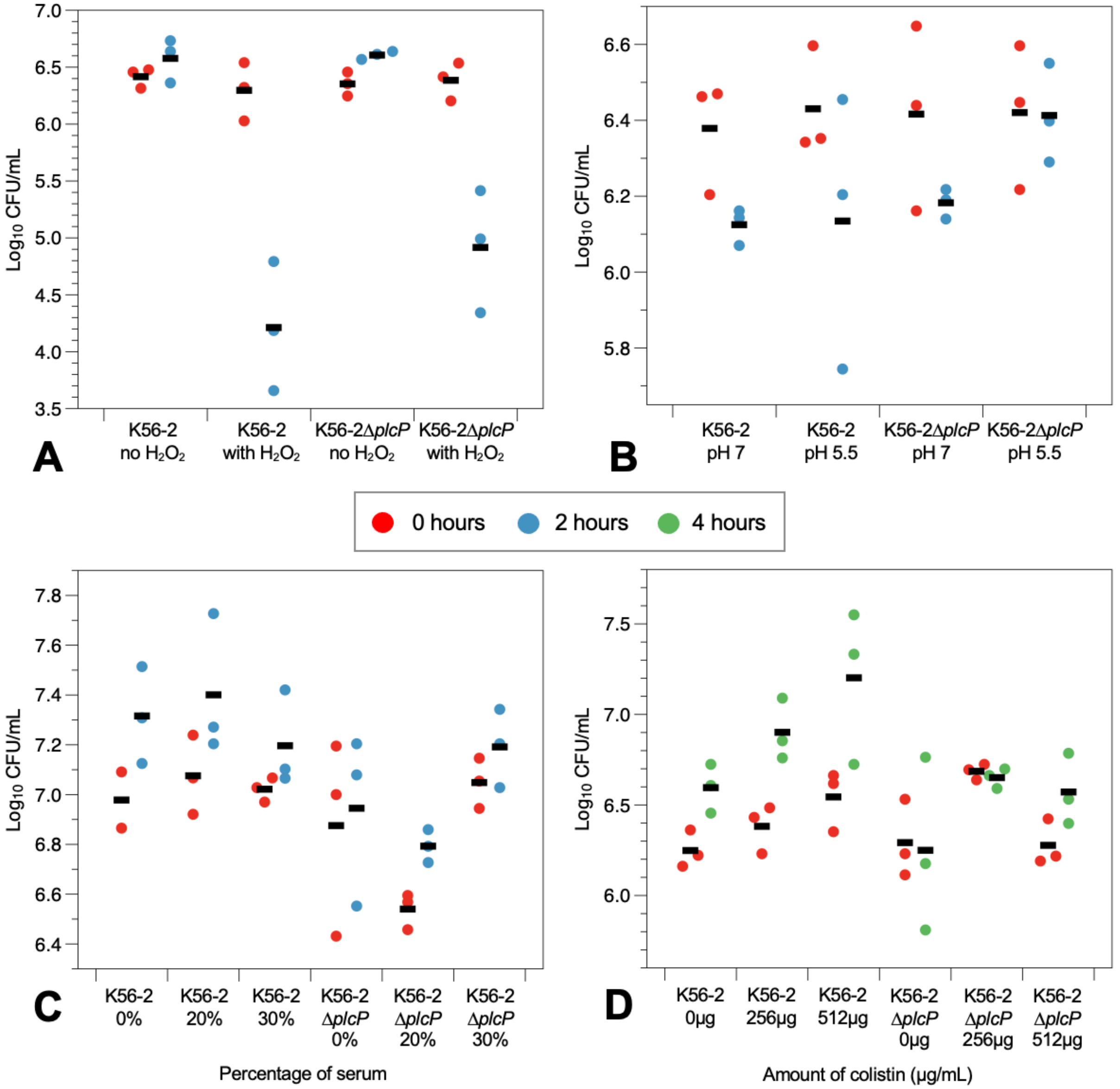


**Supplementary Figure 5. Sensitivity of wild type *B. cenocepacia* and the *plcP* mutant to various stresses.** (A) Sensitivity to $H_2O_2$ measured by changes of $log_{10}$ CFU/mL after 2 hours of exposure to 1.5 mM $H_2O_2$. (B) Sensitivity to pH measured by changes of $log_{10}$ CFU/mL after 2 hours of exposure to pH 5.5. (C) Sensitivity to human serum measured by changes of $log_{10}$ CFU/mL after 2 hours of exposure to 20% (v/v) and 30% (v/v) serum. (D) Sensitivity to antimicrobial peptides measured by changes of $log_{10}$ CFU/mL after 4 hours of exposure to 256 μg/mL and 512 μg/mL colistin.
